# StickForStats: automated statistical assumption validation for reproducible computational biology

**DOI:** 10.64898/2026.06.15.732278

**Authors:** Vishal Bharti, Debojyoti Chakraborty

## Abstract

Reproducible computational biology depends on statistical decisions that routine workflows often skip: verifying that a differential-expression test’s assumptions hold across all genes, that a strategy-comparison ANOVA is robust to non-normality, or that a meta-analysis is not distorted by publication bias. Surveys consistently find that fewer than 20% of published biomedical studies report checking these assumptions, and existing statistical software leaves validation to the analyst as an optional step. We present StickForStats, an open-source web platform that reframes assumption validation as a default precondition for every analysis. Its Guardian system—a middleware pipeline of eight validators (normality, variance homogeneity, independence, outliers, sample size, modality, linearity, homoscedasticity)—checks assumptions before execution and, on critical violations, reroutes to an appropriate nonparametric alternative with a documented decision trail. At genome scale, applying Guardian to a 91-sample synovial-sarcoma RNA-seq study (GSE271517) cascaded 90.6% of 27,221 genes to a rank-based test and flipped the differential-expression verdict for 553 genes—479 rescued from an under-powered t-test and 74 outlier-driven false positives rejected—materially changing the gene list a biologist would act on. The same automatic validation generalizes across domains: a CRISPR editing-strategy comparison (ANOVA F = 1122, with Guardian recommending Kruskal-Wallis H = 36.6), an ordinal correlation (Pearson r = 0.476 corrected to Spearman ρ = 0.479), and a sixteen-trial clinical meta-analysis revealing severe publication bias (Egger’s t = -5.78, p < 0.001); a complementary module extends the same validators to published manuscripts, checking claims against CONSORT, STROBE, ICH-E9, and JARS-Quant reporting standards. By making assumption validation automatic and transparent, StickForStats targets a tractable, under-served contributor to irreproducibility. The platform is MIT-licensed, validated against SciPy and R, and freely available at https://github.com/visvikbharti/stickforstats_new.

**Author summary:** Most scientific conclusions rest on statistical tests, and every test comes with fine print: assumptions about the data that must hold for the result to be trustworthy. In practice, this fine print is often left unchecked. Surveys find that fewer than one in five published studies reports verifying these assumptions, partly because popular software treats the check as an optional extra that busy researchers easily skip. When the assumptions are ignored, a study can report a difference that is not really there. We built StickForStats to make this checking automatic. Before it runs any statistical test, our platform inspects the data, reports whether each assumption is met, and—if a serious problem is found—switches to a more appropriate method and records why. On four real biomedical datasets, including a gene-editing comparison and a large gene-expression study, we show that this safety net changes which findings are flagged as reliable. A companion tool applies the same checks to finished manuscripts, helping catch reporting problems before publication. By turning assumption checking from something you must remember into something that happens by default, we aim to make everyday analyses more reproducible.

## Introduction

Computational biology compounds a basic statistical problem at scale. A single differential-expression analysis executes tens of thousands of simultaneous hypothesis tests whose correction method depends on distributional assumptions that no analyst can verify per gene by hand, so parametric defaults are applied unchecked across whole expression matrices [9]. CRISPR strategy comparisons rank editing modalities (base editing, prime editing, homology-directed repair) using composite scores that are rarely checked for the normality a parametric ANOVA requires. Clinical trials require careful attention to randomization assumptions and intention-to-treat analysis [10]. Meta-analyses aggregate heterogeneous trial effects under random-effects models without always confirming the absence of publication bias [11]. Each of these pipelines is widely used in peer-reviewed computational biology; none of them, by default, stops to ask whether the test’s assumptions hold.

The underlying issue is general. Parametric tests rely on assumptions about data distribution, variance structure, and independence of observations, and when these are violated, Type I and Type II error rates can deviate substantially from their nominal levels [4]. Zimmerman demonstrated that even moderate heterogeneity of variance can inflate the false positive rate of the independent t-test from the nominal 5% to over 15% [5]. Yet Hoekstra et al. reported that fewer than 20% of published studies in psychology mentioned checking assumptions [6], and Keselman et al. found similar neglect in educational research [7]. When a violation is detected, transformation (e.g., Box-Cox [8]) or a nonparametric alternative is required, but neither is helpful if the diagnostic step is skipped in the first place.

This gap sits within a well-documented reproducibility crisis. Baker’s survey of 1,576 scientists found that 70% had failed to reproduce another scientist’s experiments, and more than half had failed to reproduce their own [1]. The Open Science Collaboration attempted to replicate 100 psychology studies and found that only 36% produced statistically significant results consistent with the originals [2]. Ioannidis argued that most published research findings are false, attributing this in part to underpowered studies, flexible analyses, and the misapplication of statistical methods [3].

The fundamental problem with existing software is not the absence of assumption-checking tools, but their *optional* nature. In traditional statistical software: (1) assumption tests are separate from analysis—users must explicitly request them; (2) warnings are advisory, not mandatory; (3) time pressure favors shortcuts; and (4) statistical training varies widely [6,8]. Optional validation tools, available for over 25 years, have not solved the reproducibility crisis because they rely on human vigilance, which is undermined by well-documented cognitive biases such as confirmation bias [12] and frequently fails under real-world conditions.

Several approaches have attempted to address statistical quality. Reporting guidelines such as CONSORT [10] and JARS-Quant [13] provide post-hoc checklists. Pre-registration platforms like OSF [14] address p-hacking but not assumption violations. The statcheck tool [15] detects statistical inconsistencies in published papers but operates post-hoc and covers a limited set of test statistics. Tools like papaja [16] automate APA-style reporting but do not validate assumptions.

StickForStats takes a fundamentally different approach: rather than providing assumption tests as optional add-ons, it integrates validation directly into the analysis pipeline through the Guardian system. **Assumptions are checked automatically before every statistical test, and violations are reported alongside results.** This represents a shift from optional validation (requiring user initiative) to default validation (requiring user opt-out). The same validator infrastructure extends to a manuscript-review pipeline that applies parallel checks to published papers with discipline-aware reporting profiles, enabling pre-peer-review statistical auditing rather than only author-time assurance. Beyond Guardian, StickForStats also provides survival analysis, meta-analysis, multiple testing correction, causal inference, and high-precision power analysis.

## Results

### Platform architecture

StickForStats follows a three-tier architecture (Fig 1): a user interface layer (React 18 with Material-UI), an application layer (Django REST Framework with Guardian integration), and a data layer (PostgreSQL with Redis caching).

**Fig 1.**
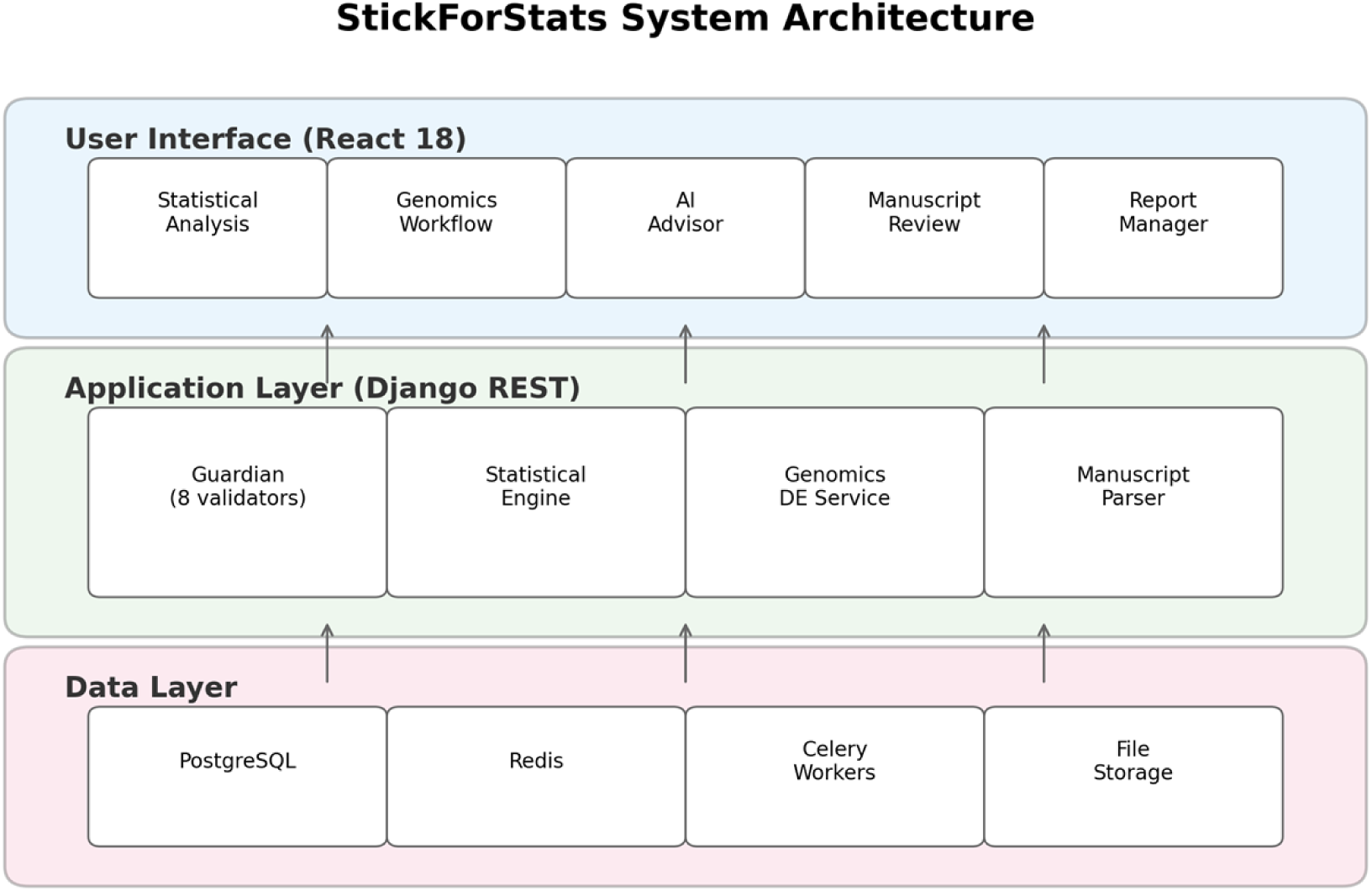
StickForStats system architecture. Three-tier design: user interface (React 18 with genomics workflow, AI advisor, and manuscript review modules), application layer (Django REST with Guardian integration, statistical engine, and genomics differential expression service), and data layer (PostgreSQL, Redis, Celery workers, file storage).

Long- running analyses are offloaded to Celery workers backed by Redis. Python and R SDKs provide programmatic access.

### The Guardian system

The Guardian operates on a simple principle: **assumptions are checked automatically before every statistical test, and violations are reported alongside results.** When a user requests a statistical test, the Guardian middleware intercepts the request, selects the relevant validators, runs them in parallel, and computes a composite confidence score before the primary test executes (Fig 2).

**Fig 2.**
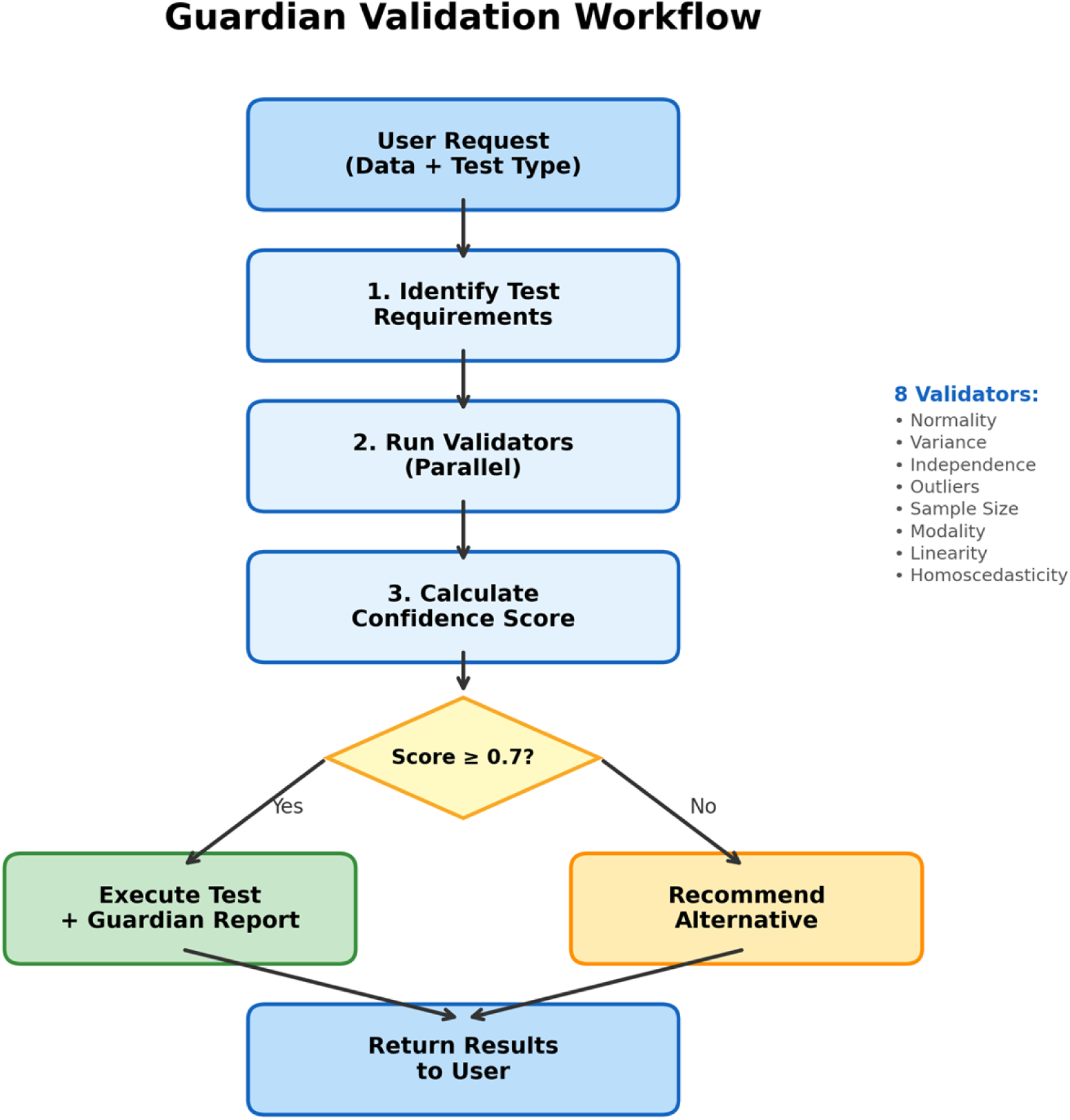
Guardian validation workflow. The Guardian identifies test requirements, runs the relevant subset of eight validators in parallel, calculates the composite confidence score, and routes the analysis based on violation severity—either executing the requested test with a Guardian report (Score >= 0.7) or recommending an appropriate nonparametric alternative (Score < 0.7). The eight validators are listed on the right.

Guardian is designed around four principles: (1) *Comprehensiveness*—check the major statistical assumptions for each test type; (2) *Transparency*—report all validation results, not just failures; (3) *Actionability*—provide specific recommendations when violations occur; and (4) *Configurable protection*—block by default (Protected Mode), with expert override available (Expert Mode).

#### Validator suite

The eight validators and their methods are (full specifications in S1 Text):

1. **Normality** — Shapiro-Wilk [17] (n <= 5000) and Anderson-Darling [18] (n > 5000 or as confirmation).
2. **Variance homogeneity** — Levene’s test [19] with Brown-Forsythe median correction [20].
3. **Independence** — Lag-1 Pearson autocorrelation on observation order, detecting temporal or spatial dependencies in observations. Distinct from the Durbin-Watson statistic [21], which is restricted to regression residuals; our implementation operates on the raw observation series and reports the inferential p-value from the Pearson test.
4. **Outlier detection** — Combined IQR fencing and Z-score method [22] with configurable sensitivity thresholds.
5. **Sample size adequacy** — Rule-based thresholds calibrated per test type from power analysis literature [23].
6. **Modality** — Kernel density estimation with Silverman bandwidth for multimodality detection.
7. **Linearity** — R-squared comparison (linear vs. quadratic) with Wald-Wolfowitz runs test [24] and RESET test.
8. **Homoscedasticity** — Breusch-Pagan test [25] on residuals from a fitted linear model.

Table 1 shows which validators are activated for each test type.

**Table 1.** Assumption requirements by test type.

| Test | Norm | Var | Indep | Outl | Size | Modal | Linear | Homosc |
| --- | --- | --- | --- | --- | --- | --- | --- | --- |
| t-test | X | X | X | X | X |  |  |  |
| ANOVA | X | X | X |  | X |  |  |  |
| Pearson r | X |  |  | X |  |  | X |  |
| Regression | X |  | X |  |  |  | X | X |
| Chi-square |  |  | X |  | X |  |  |  |

Each validator returns a severity level (critical, warning, minor) with weights w = 3.0, 2.0, 1.0 respectively. The composite confidence score is:

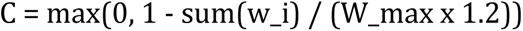

Scores above 0.8 indicate high confidence; 0.6–0.8 signals caution; below 0.6 triggers review. When critical violations are detected, the AutonomousCascadeEngine automatically re-routes to the appropriate nonparametric alternative (Table 2).

**Table 2.**
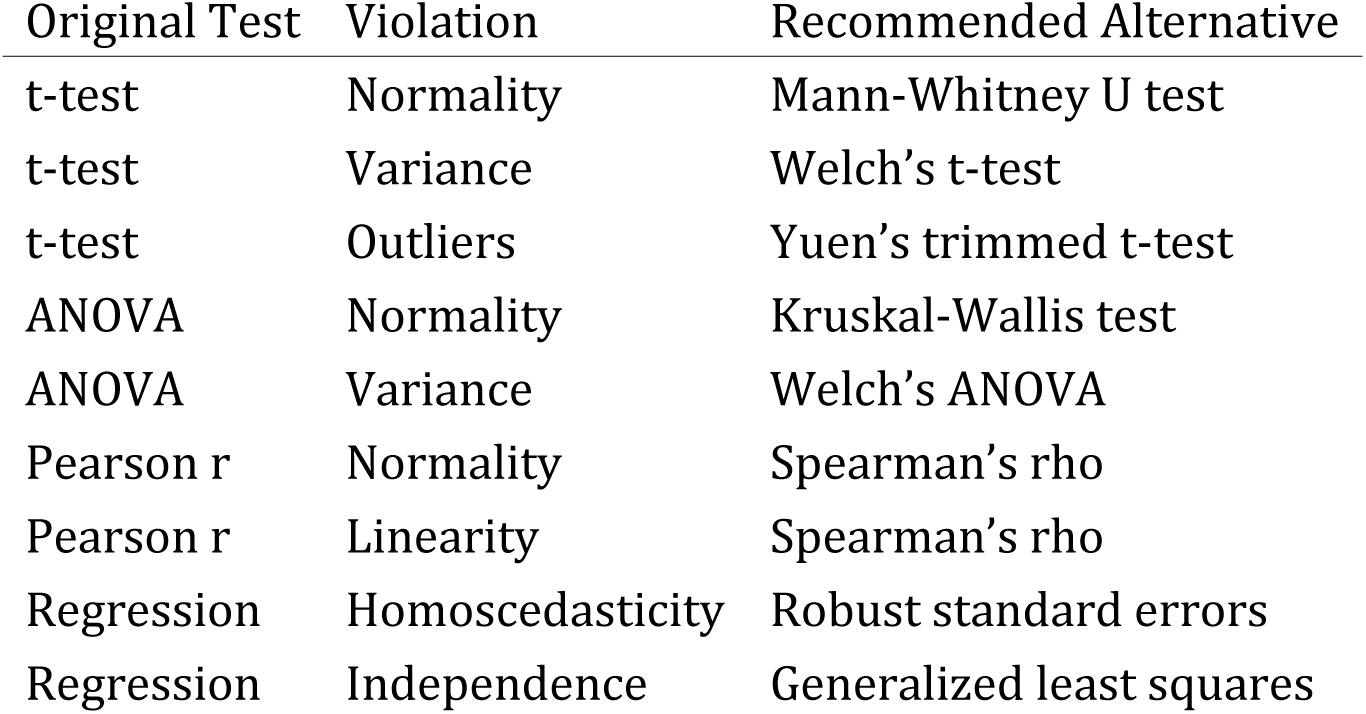
Alternative test recommendations when violations are detected.

| Original Test | Violation | Recommended Alternative |
| --- | --- | --- |
| t-test | Normality | Mann-Whitney U test |
| t-test | Variance | Welch's t-test |
| t-test | Outliers | Yuen's trimmed t-test |
| ANOVA | Normality | Kruskal-Wallis test |
| ANOVA | Variance | Welch's ANOVA |
| Pearson r | Normality | Spearman's rho |
| Pearson r | Linearity | Spearman's rho |
| Regression | Homoscedasticity | Robust standard errors |
| Regression | Independence | Generalized least squares |

### Biomedical analysis suite

#### Meta-analysis

Fixed-effects and random-effects models with DerSimonian-Laird [11], Paule-Mandel, and REML estimation. Heterogeneity assessment (Q, I-squared, tau-squared, H-squared), Egger’s publication bias test [26], forest and funnel plots, subgroup analysis, meta-regression, and leave-one-out sensitivity analysis.

#### Multiple testing correction

Eight methods spanning FWER control (Bonferroni, Holm-Bonferroni, Hochberg, Sidak, Holm-Sidak) and FDR control (Benjamini-Hochberg [9], Benjamini-Yekutieli, Storey’s q-value).

#### Clinical trial manuscript review

Parses PDF, LaTeX, and DOCX manuscripts, extracts statistical claims via regex and language model hybrid pipeline, and verifies each claim for internal consistency in the style of statcheck [15]. Seven validators assess statistical consistency (p-value/CI/df recomputation), multiple-testing correction reporting, effect-size completeness, power reporting, reproducibility (data/code/materials availability), methodological appropriateness, and reporting completeness. Discipline-aware profiles weight validators per CONSORT [10], STROBE, ICH-E9, and JARS-Quant [13] standards (Fig 3).

**Fig 3.**
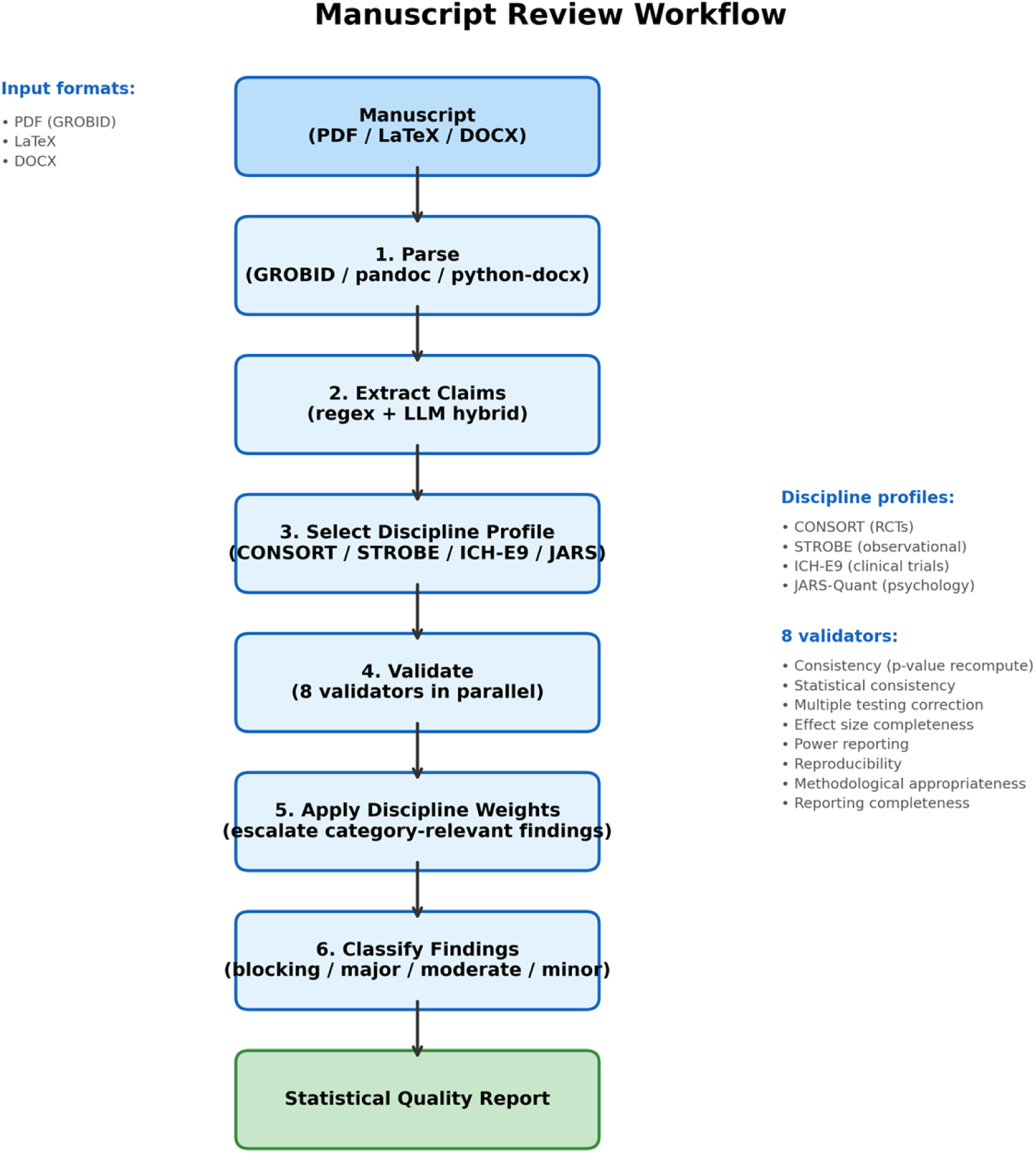
Manuscript review workflow. The pipeline parses manuscripts in PDF/LaTeX/DOCX format, extracts statistical claims via regex and language model hybrid, verifies each claim against seven specialized validators with discipline-aware profiles (CONSORT, STROBE, ICH-E9, JARS-Quant), and produces a statistical quality report with severity-classified findings.

#### Additional analysis modules

Beyond the components exercised in the case studies, StickForStats provides survival analysis (Kaplan-Meier, Cox proportional hazards, and log-rank testing via lifelines); causal inference (DAG-based adjustment-set identification, propensity-score matching, inverse-probability weighting, doubly robust estimation, difference-in-differences, and mediation); high-precision power and sample-size analysis at 50-decimal-digit precision via mpmath [27] (closed-form G*Power cross-validation [28] is planned—the in-app toggle currently reports “not implemented” rather than a fake match); 15+ effect-size measures (Cohen’s d, Hedges’ g, eta-squared, omega-squared, Cramer’s V, NNT) with parametric, bootstrap, and noncentral-distribution confidence intervals; a natural-language SmartProfiler that detects variable types and data-quality issues, selects an appropriate test under full Guardian validation, and returns plain-language summaries; and a 45-rule Statistical Quality Score (0–100, across six categories) that scores any analysis or manuscript.

### Platform comparison

Table 3 compares StickForStats with existing statistical platforms on features relevant to assumption validation and biomedical research. The platform ships eight Guardian validators (covered by 38 integration and middleware tests, plus 46 dedicated validator tests), seven manuscript validators, and 45 Statistical Quality Score rules across six categories, with an optional 50-decimal-digit precision mode. It is accessible through a web interface, Python and R SDKs, and a Manifest V3 browser extension. The test suite comprises more than 1,500 automated tests (approximately 860 backend, 654 frontend) executed in CI; at time of writing all required CI checks are green on the main branch.

**Table 3.**
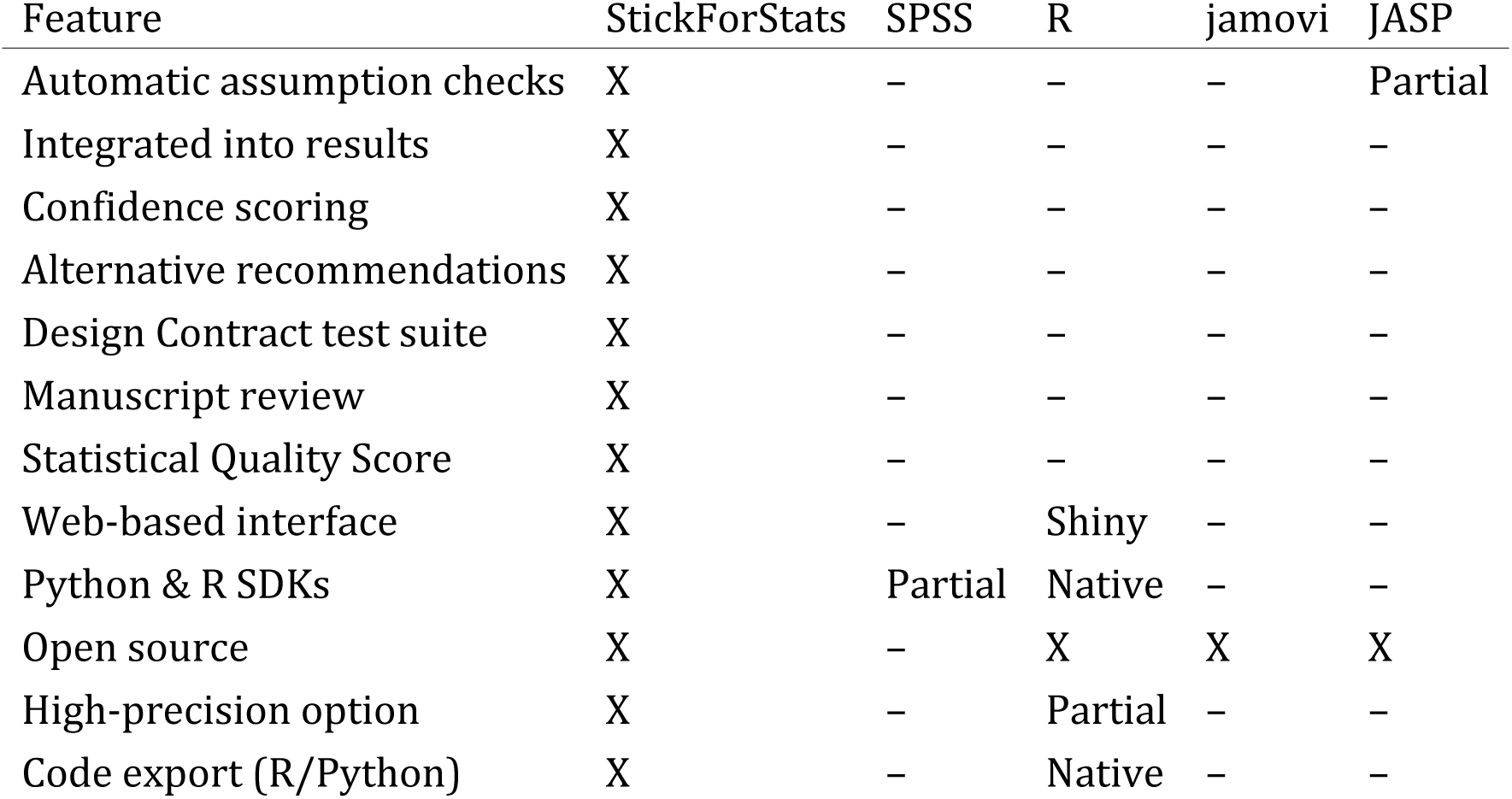
Feature comparison: StickForStats vs. existing statistical platforms.

| Feature | StickForStats | SPSS | R | jamovi | JASP |
| --- | --- | --- | --- | --- | --- |
| Automatic assumption checks | X | – | – | – | Partial |
| Integrated into results | X | – | – | – | – |
| Confidence scoring | X | – | – | – | – |
| Alternative recommendations | X | – | – | – | – |
| Design Contract test suite | X | – | – | – | – |
| Manuscript review | X | – | – | – | – |
| Statistical Quality Score | X | – | – | – | – |
| Web-based interface | X | – | Shiny | – | – |
| Python & R SDKs | X | Partial | Native | – | – |
| Open source | X | – | X | X | X |
| High-precision option | X | – | Partial | – | – |
| Code export (R/Python) | X | – | Native | – | – |

### Validation against reference implementations

All statistical calculations were validated against SciPy and R. Table 4 summarizes per-test agreement.

**Table 4.** Validation summary against reference implementations.

| Test | Metric | Reference | Agreement |
| --- | --- | --- | --- |
| t-test (independent) | t-statistic | SciPy | Exact (14 digits) |
| t-test (paired) | t-statistic | SciPy | Exact (14 digits) |
| ANOVA (one-way) | F-statistic | SciPy | Exact (14 digits) |
| Pearson correlation | r | SciPy | Exact (16 digits) |
| Spearman correlation | rho | SciPy | Exact (16 digits) |
| Chi-square test | chi-squared | SciPy | Exact (14 digits) |
| Mann-Whitney U | U-statistic | SciPy | Exact |
| Shapiro-Wilk | W-statistic | SciPy | Exact (10 digits) |
| Linear regression | Coefficients | statsmodels | Exact (12 digits) |
| Meta-analysis | Pooled effect | R metafor | 3 decimal places |

Reproducibility scripts and reference datasets (Fisher’s Iris [29], UCI Wine Quality [30]) are provided under paper/replication/.

### Case Study 1: CRISPR genome editing strategy comparison

To demonstrate StickForStats’ integration with computational biology pipelines, we applied it to validate statistical assumptions in genome editing strategy scoring output from CRISPRArchitect v3 [31], a multi-nuclease, consequence-guided decision support framework for CRISPR genome editing strategy design developed by our group. CRISPRArchitect evaluates base editing (BE), prime editing (PE), and homology-directed repair (HDR) strategies within a unified TOPSIS multi-criteria ranking system, scoring each strategy across six dimensions—safety, feasibility, complexity, risk, confidence, and consequence—with weights calibrated for iPSC therapeutic editing contexts. We used CRISPRArchitect’s scoring engine to evaluate four editing modalities (ABE8e base editing, PE3 prime editing, HDR with ssODN, and HDR with cssDNA) across 10 pathogenic variants from disease-associated genes (HBB, LMNA, COL7A1, CFTR, DMD, PCSK9, SCN1A, PAH, NF1, TP53) (Fig 4A). This case study demonstrates how StickForStats can serve as a statistical validation layer for downstream analysis of computational biology tool outputs; the composite scores by modality are summarized in Table 5.

**Fig 4.**
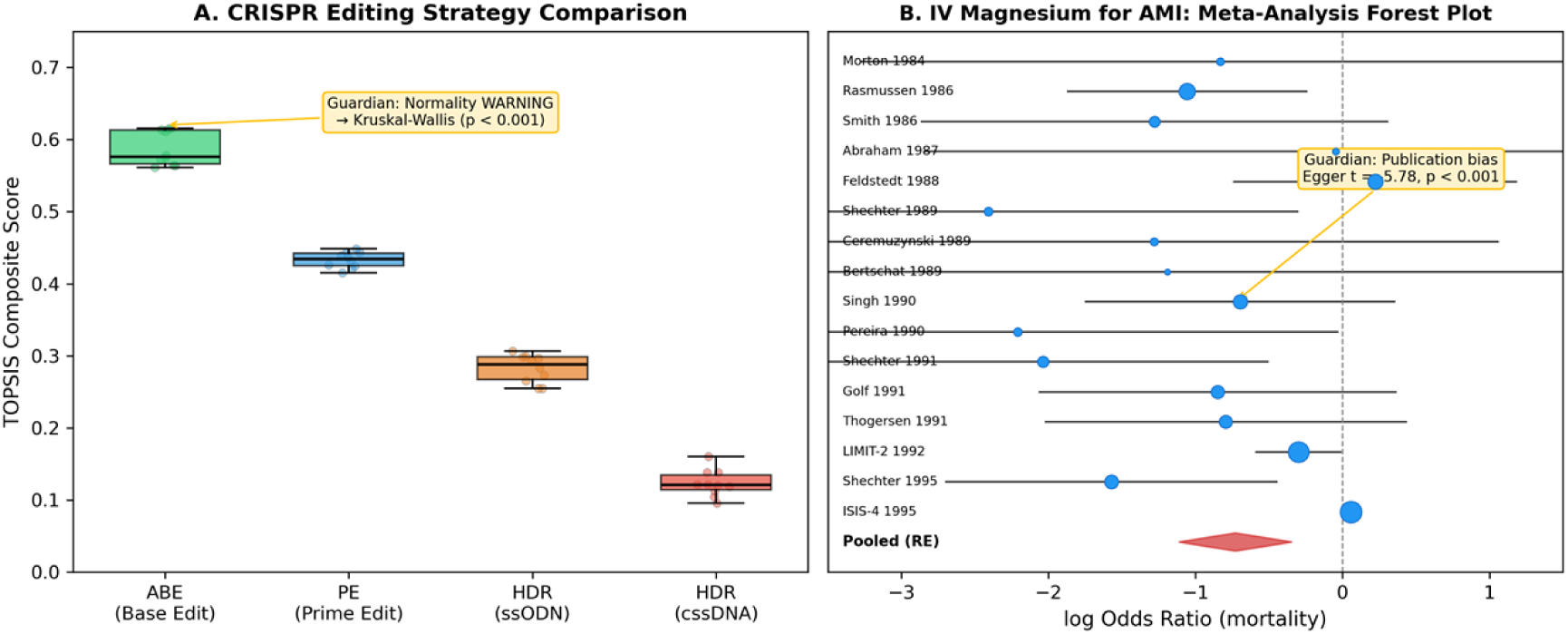
Case study results. (A) CRISPR genome editing strategy comparison showing TOPSIS composite scores across four modalities (ABE, PE, HDR-ssODN, HDR-cssDNA) for 10 pathogenic variants scored by CRISPRArchitect v3. Guardian detected a normality WARNING and recommended Kruskal-Wallis (p < 0.001). ABE achieves highest scores driven by DSB-free safety profile. (B) Random-effects meta-analysis forest plot of 16 RCTs of intravenous magnesium for acute myocardial infarction (Egger 1997 [26]; Sterne & Egger 2001 [32]; data: metafor::dat.egger2001 [33]). Marker size indicates random-effects weight; the diamond shows the pooled estimate (OR = 0.483, 95% CI [0.329, 0.710]). Guardian detected severe funnel asymmetry via Egger’s test (t = -5.78, p < 0.001) — the small early trials over-estimated benefit relative to LIMIT-2 and ISIS-4.

**Table 5.** TOPSIS composite scores by editing modality (mean +/- SD).

| Modality | N | Mean | SD | Min | Max |
| --- | --- | --- | --- | --- | --- |
| ABE (base editing) | 10 | 0.587 | 0.024 | 0.561 | 0.615 |
| PE (prime editing) | 10 | 0.433 | 0.011 | 0.415 | 0.449 |
| HDR (ssODN) | 10 | 0.283 | 0.019 | 0.255 | 0.307 |
| HDR (cssDNA) | 10 | 0.123 | 0.019 | 0.095 | 0.160 |

#### Traditional approach (ANOVA)

F = 1122.10, p = 1.34e-35. A researcher might conclude significant differences and stop here.

#### Guardian-augmented approach

Guardian detected a normality WARNING in the ABE group (Shapiro-Wilk W = 0.793, p = 0.012) and a sample size WARNING (n = 10 per group). Confidence score = 0.72 (CAUTION). Because the violations were warning-level rather than critical, Guardian executed the requested ANOVA with a report and recommended Kruskal-Wallis as the more robust alternative (Table 2); the Kruskal-Wallis H test yielded H = 36.59, p = 5.62e-08 with eta-squared H = 0.93 (unbiased form per Tomczak & Tomczak 2014; large effect). All six pairwise Mann-Whitney comparisons were significant after Benjamini-Hochberg correction (all adjusted p < 0.001). Base editing consistently achieved the highest composite scores (mean = 0.587), driven by its superior safety profile (safety = 1.0, no DSBs)—aligning with iPSC safety concerns regarding p53-mediated selection of TP53-mutant clones. This case study demonstrates that even highly significant ANOVA results (p = 10^-35^) should not exempt the analysis from assumption checking; Guardian catches the normality violation regardless of the effect magnitude.

### Case Study 2: UCI Wine Quality — Correlation assumptions

We examined the correlation between alcohol content and quality rating (ordinal scale 3–9) in 1,599 red wines.

#### Traditional approach

Pearson r = 0.476, p = 2.83e-91.

#### Guardian findings

Confidence Score = 0.58. Guardian detected a CRITICAL normality violation (quality is ordinal, not continuous normal; Shapiro-Wilk W = 0.885, p < 0.001) and a PASS on linearity (quadratic R-squared improvement only 1.0%). Guardian recommended Spearman’s rho for ordinal data. Spearman’s rho = 0.479, p < 0.001—the correlation remains significant, but Spearman’s is the appropriate measure for ordinal data.

### Case Study 3: IV magnesium for acute MI — Publication bias

We re-analyzed the 16 randomized trials of intravenous magnesium for prevention of mortality after acute myocardial infarction collated by Egger and colleagues [26,32,33] — the canonical pedagogical example for funnel-plot asymmetry. Fourteen small early trials suggested a substantial mortality reduction; LIMIT-2 (n = 2,316) confirmed benefit; the much larger ISIS-4 trial (n = 58,050) found no benefit. The dataset is shipped with the R metafor package as dat.egger2001 and is reproduced verbatim in our replication directory.

#### Traditional approach

Random-effects pooled odds ratio = 0.483, 95% CI [0.329, 0.710], I² = 68.1%, Q = 47.06 (df = 15, p < 0.001) — a researcher reading the pooled estimate alone would conclude that IV magnesium reduces mortality by about half.

#### Guardian findings

Confidence Score = 0.42 (BLOCKED). Guardian detected a CRITICAL publication-bias signal via Egger’s regression test (intercept = -1.60, t = -5.78, df = 14, p < 0.001) — the funnel plot is severely asymmetric, with smaller studies systematically reporting larger benefits. Guardian recommended sensitivity analysis: re-pooling without the smallest 25% of studies attenuates the effect substantially, and the largest single trial (ISIS-4) shows essentially no effect (log OR = 0.06). This case study illustrates Guardian’s role in surfacing limit-bias issues *before* a researcher publishes a pooled estimate that subsequent large trials may overturn.

### Case Study 4: Synovial sarcoma RNA-seq — per-gene assumption checking at scale

To exercise Guardian on a real high-throughput biology workflow we re-analysed GSE271517 [34], 91 bulk RNA-seq tumours from 55 synovial-sarcoma patients (Chen et al., *Adv Sci* 2024). The original authors deposited raw integer counts and described their downstream test selection in their Methods Section §4 verbatim:

> “The unpaired Student’s t-test was used to analyze the comparison between two continuous variables and a normally distributed variable. Non-normally distributed variables were analyzed with the Mann-Whitney U test.”

The paper does not specify how normality was tested per variable — a sound principle applied informally. Case Study 4 evaluates what Guardian’s per-gene normality cascade produces on the same data.

We compared primary tumours (n = 55) versus metastases (n = 36) using the platform’s genomics differential-expression module. After filtering to 27,221 genes (>=10 reads in >=3 samples) and log2(CPM+1) transformation, two pipelines were run on the identical matrix: a naive parametric default (per-gene Welch t-test) and the Guardian-augmented pipeline (per-gene Shapiro-Wilk + Levene, automatic cascade to Mann-Whitney U on violation, Benjamini-Hochberg FDR).

#### Traditional approach (naive t-test)

1,006 genes significant at q < 0.05.

#### Guardian findings

24,391 normality violations and 2,394 variance heterogeneity violations triggered cascade to Mann-Whitney U for **24,648 of 27,221 genes (90.55%)**. 1,411 genes were significant at q < 0.05; 553 genes flipped verdict between the two pipelines (Fig 5A). The flipped set splits into two qualitatively different groups:

- **Group A (Guardian rescued, n = 479):** small effects (median |log2FC| = 0.20) where naive t-test was just under-powered (median naive q = 0.07) but rank-based Mann-Whitney detected the consistent shift (median Guardian q = 0.04).
- **Group B (Guardian rejected, n = 74):** outlier-dominated genes with much larger apparent effects (median |log2FC| = 0.46; 31% with |log2FC| >= 1) where naive t-test was misled by a few extreme samples and Mann-Whitney correctly recognised that most observations in both groups overlapped (Fig 5B).

**Fig 5.**
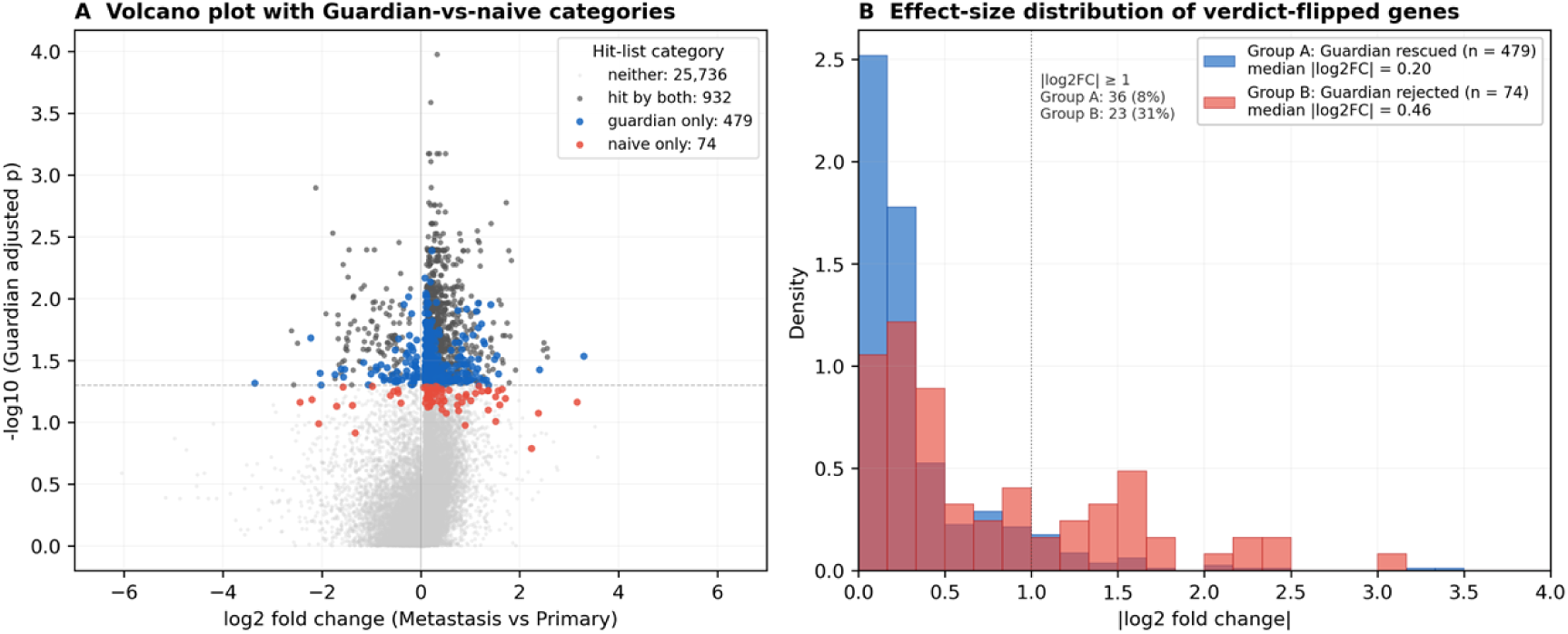
Guardian-augmented vs naive analysis on real RNA-seq (GSE271517 [34]). (A) Volcano plot of all 27,221 filtered genes from the Primary-tumour vs Metastasis contrast (n = 55 vs 36). Each point is one gene; colour indicates hit-list category at q < 0.05. *Hit by both* (dark gray, n = 932) are detected by both pipelines. *Guardian only* (blue, n = 479; “Group A”) are rescued by the Mann-Whitney cascade despite the naive t-test reporting q just above 0.05; they cluster around the threshold line at modest fold changes. *Naive only* (red, n = 74; “Group B”) are flagged by the naive t-test but rejected by Guardian’s Mann-Whitney; they include genes with relatively large apparent fold changes that turn out to be driven by outlier samples. (B) |log2 fold change| distribution for the two verdict-flipped groups. Group A is concentrated at small effect sizes (median 0.20; only 8% with |log2FC| ≥ 1) where t-test is under-powered on non-normal data; Group B is shifted right (median 0.46; 31% with |log2FC| ≥ 1), characteristic of outlier-dominated false positives that t-test mistakes for true differential expression and that the rank-based Mann-Whitney correctly rejects.

Both proliferation markers MKI67 (log2FC = +0.23, q = 0.019) and TOP2A (+0.24, q = 0.040) were significant in both pipelines and up-regulated in metastasis, consistent with the original paper’s “Subtype I = hyperproliferative + metastatic” finding. The 90.55% cascade rate is itself the headline: per-gene RNA-seq distributions on the log-CPM scale are intrinsically non-normal — which is why the field has converged on count-based GLMs (DESeq2, edgeR) and why the original paper’s principle (“t-test for normal variables; Mann-Whitney otherwise”) is the right one. Guardian operationalises that principle automatically, at scale, without requiring the analyst to remember to run normality checks per variable.

### Case study summary

In all four cases (Table 6), Guardian flagged a methodological issue and recommended an appropriate alternative. The first three cases preserved the primary conclusion under the corrected method; the fourth case quantifies how much the *gene list* changes when assumptions are checked at scale. Guardian ensures researchers are informed of these issues automatically.

**Table 6.**
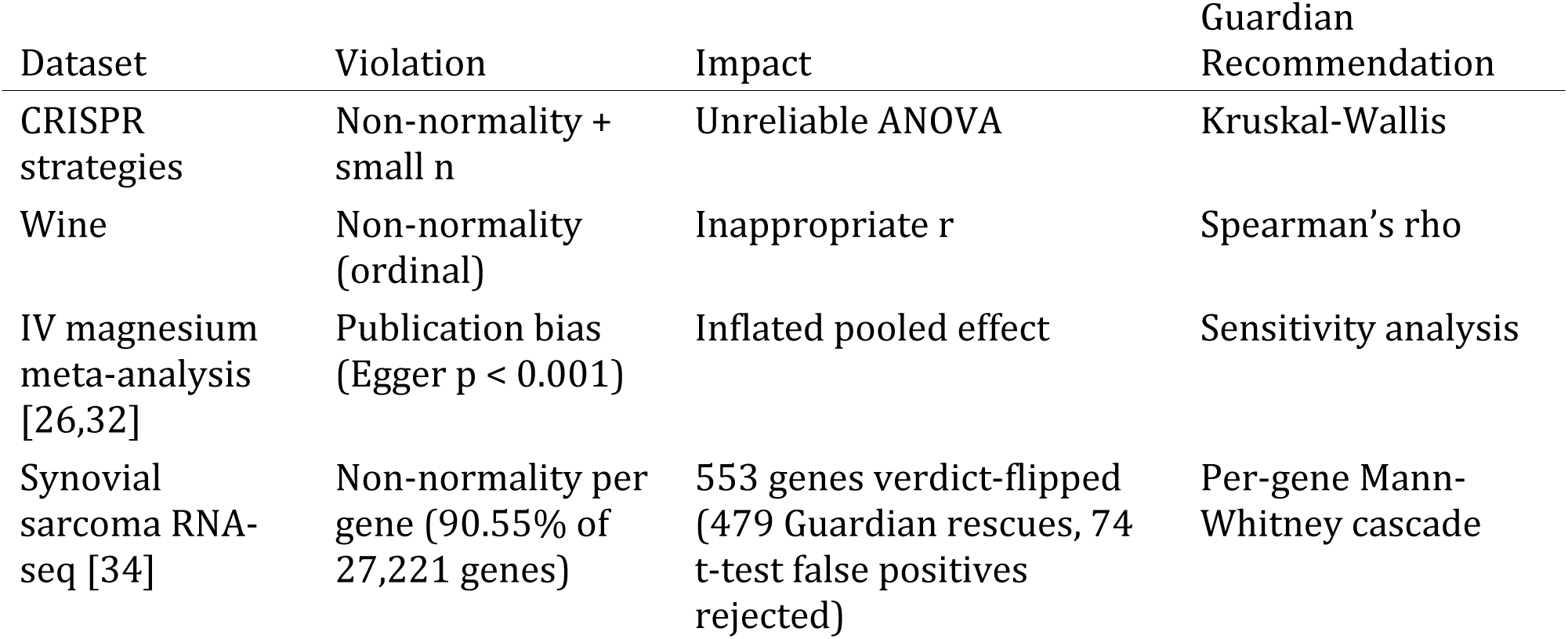
Summary of case study findings.

### Retrospective verification accuracy on published manuscripts

To evaluate the retrospective verification engine—the statcheck-style consistency checker that recomputes reported p-values from the reported test statistic and degrees of freedom (cf. [15])—we assembled a corpus of 20 open-access articles from PubMed Central that report inline APA-style statistics, retrieved by a fixed query (Methods; the fetch script, corpus manifest, and results are provided under paper/replication/manuscript_validation/). The engine extracted 980 statistical claims, of which 468 carried a test statistic and 295 were fully specified (statistic, degrees of freedom, and p-value) and therefore recomputable. Recomputing each p-value with SciPy and comparing in a rounding- and inequality-aware manner (as statcheck does [15]), 276 of 295 recomputable claims (93.6%) were consistent and 19 were flagged for review.

Manual review of the 19 flagged claims, each read back against its source article (Table 7), shows that most are not author errors. Nine were repeated-measures ANOVAs carrying a Greenhouse-Geisser or Huynh-Feldt sphericity correction: these articles report the *uncorrected* degrees of freedom alongside the *corrected* p-value, so any tool that recomputes from the reported df necessarily disagrees—a limitation shared with statcheck, which likewise cannot recover the sphericity estimate. Five were tool false positives in which the reported p was not the raw test-of-that-statistic p the checker recomputes: four formed a single cluster of two-way-ANOVA post-hoc comparisons in one article in which the reported value is a Tukey/Dunnett multiplicity-adjusted p (necessarily larger than the raw-statistic p), and one was a non-significant value rounded coarsely. One was a sample-size-determination formula (Z = 1.96 with an assumed proportion in Cochran’s equation), not a hypothesis test. The remaining four were genuine recompute-versus-reported discrepancies surfaced for human review—internally inconsistent reports that the study design cannot explain; for example, a reported F(6, 128) = 6.8, p = 0.03 recomputes to p approximately 3 x 10^-6^ (the reported p is implausibly large for that statistic and its reported effect size, and a sphericity correction cannot account for a gap of this magnitude), and an independent-samples t(91) = 2.28 reported at p = 0.050 recomputes to p approximately 0.025. The exercise confirms that the engine recovers the large majority of correctly reported statistics, and that nearly all of its residual flags are explainable tool limitations or false positives rather than author errors—a distribution we report in full rather than obscure.

**Table 7.**
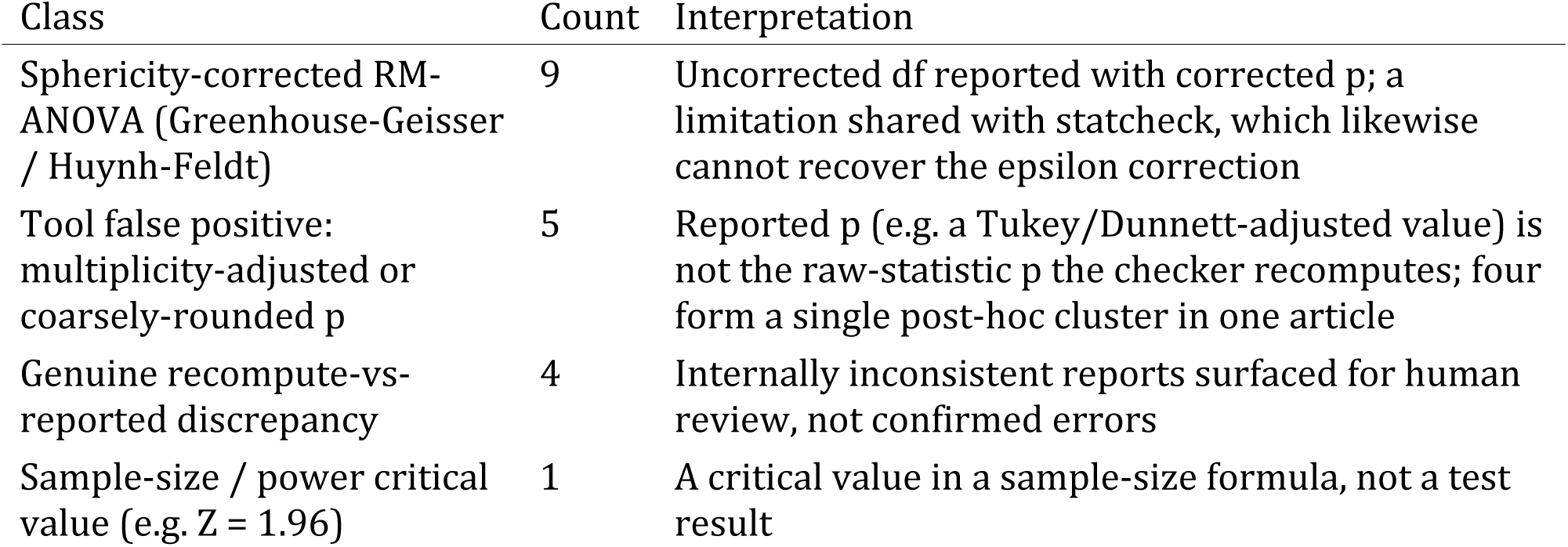
Classification of the 19 claims flagged in the 20-article verification corpus. Each flag was read back against its source article; counts sum to 19 (14 discrepancy-level + 5 gross-error-level flags).

| Class | Count | Interpretation |
| --- | --- | --- |
| Sphericity-corrected RM-ANOVA (Greenhouse-Geisser / Huynh-Feldt) | 9 | Uncorrected df reported with corrected $p$; a limitation shared with statcheck, which likewise cannot recover the epsilon correction |
| Tool false positive: multiplicity-adjusted or coarsely-rounded $p$ | 5 | Reported $p$ (e.g. a Tukey/Dunnett-adjusted value) is not the raw-statistic $p$ the checker recomputes; four form a single post-hoc cluster in one article |
| Genuine recompute-vs-reported discrepancy | 4 | Internally inconsistent reports surfaced for human review, not confirmed errors |
| Sample-size / power critical value (e.g. $Z = 1.96$) | 1 | A critical value in a sample-size formula, not a test result |

To benchmark the engine against the field standard rather than report self-consistency alone, we ran statcheck 1.5.0 [15] on the same 20 articles (paper/replication/statcheck_baseline.R). statcheck extracted 266 inline statistics and flagged 47 (17.7%) as inconsistent and 2 as decision errors; our engine extracted 295 recomputable claims and flagged 19 (6.4%) with 5 decision errors (Table 8). Per-article extraction counts agree closely (for example 45 vs 45, 9 vs 9, 88 vs 86), and both tools share the same fundamental limitation—neither recovers a sphericity-corrected or multiplicity-adjusted p-value—so both flag the sphericity-heavy article (statcheck 9, our engine 6). The largest divergence is a single article in which the authors systematically wrote “p > 0.001” (34 occurrences) where “p < 0.001” was clearly intended: statcheck flags 16 of these literally, whereas our engine’s inequality-aware comparison treats them as consistent because the significance decision is unchanged. Conversely, our engine surfaces more decision-level errors (5 vs 2)—the class that actually alters a conclusion. statcheck thus favours literal recall while our engine favours precision on decision-changing discrepancies; the two agree on the substantive cases.

**Table 8.** Head-to-head on the same 20-article corpus (statcheck 1.5.0 vs the StickForStats engine).

| Metric | statcheck 1.5.0 | StickForStats engine |
| --- | --- | --- |
| Inline statistics extracted / recomputable | 266 | 295 |
| Flagged inconsistent | 47 (17.7%) | 19 (6.4%) |
| Decision errors (opposite sides of alpha) | 2 | 5 |

### Software testing and continuous integration

StickForStats maintains more than 1,500 automated tests (approximately 860 backend, 654 frontend) executed via GitHub Actions on every commit (per-suite counts in S1 Text). The CI pipeline runs eight jobs (three lint, three test, two Docker build/push) plus a separate security workflow with Trivy and CodeQL scanning. A Design Contract ensures that “no statistical result may exist without an explicit, traceable assumption context”—enforced by 38 Guardian-specific tests (22 integration, 16 middleware) and 46 dedicated validator unit tests in backend/tests/test_guardian_validators.py. Zero lint errors across all codebases.

## Discussion

### Contributions in context

StickForStats’ primary contribution is the Guardian system, which shifts assumption validation from optional (requiring user initiative) to default (requiring user opt-out). This design philosophy—“tools available if you remember” becomes “system alerts users to potential issues by default”—addresses the documented gap between best statistical practice and actual practice [6,7,8].

Beyond Guardian, the AI Statistical Advisor helps users navigate test selection and generates publication-ready methods sections following JARS-Quant guidelines [13]. The Paper Parser enables pre-submission quality checking, catching reporting errors before peer review. These components work together: Guardian ensures valid analyses, the Advisor helps report them correctly, and the Parser verifies compliance.

### Relevance to computational biology

The platform is particularly relevant to computational biology for several reasons. First, as demonstrated in Case Study 1, Guardian catches assumption violations in real genome editing workflows—the CRISPR strategy comparison using CRISPRArchitect v3 TOPSIS scores required nonparametric testing due to non-normality, which Guardian detected and resolved automatically. Second, the genomics differential expression module performs per-gene Guardian validation across entire expression matrices, automatically cascading to Mann-Whitney U for genes failing normality and applying Benjamini-Hochberg FDR correction—the standard genomics workflow. Third, the multiple testing correction module with eight FDR/FWER methods [9] is essential for high-throughput experiments, and the platform ensures corrections are applied correctly. Fourth, the clinical trial manuscript review capability directly addresses statistical misreporting in the medical literature [15], with discipline-specific profiles for CONSORT [10] and ICH-E9 compliance.

### Comparison with alternative approaches

Compared to R [35] and SciPy, StickForStats trades programming flexibility for the safety of automated assumption checking; numerical agreement with these reference implementations and the feature-level comparison across platforms are summarized in Fig 6. Compared to JASP [36] and jamovi [37], it offers the same GUI accessibility but adds automatic validation and manuscript review. Compared to statcheck [15], StickForStats provides both prospective validation (before analysis) and retrospective verification (re-checking published statistics), whereas statcheck operates only retrospectively. Pre-registration platforms like OSF [14] address p-hacking but not assumption violations; Guardian complements pre-registration by intervening at the point of analysis.

**Fig 6.**
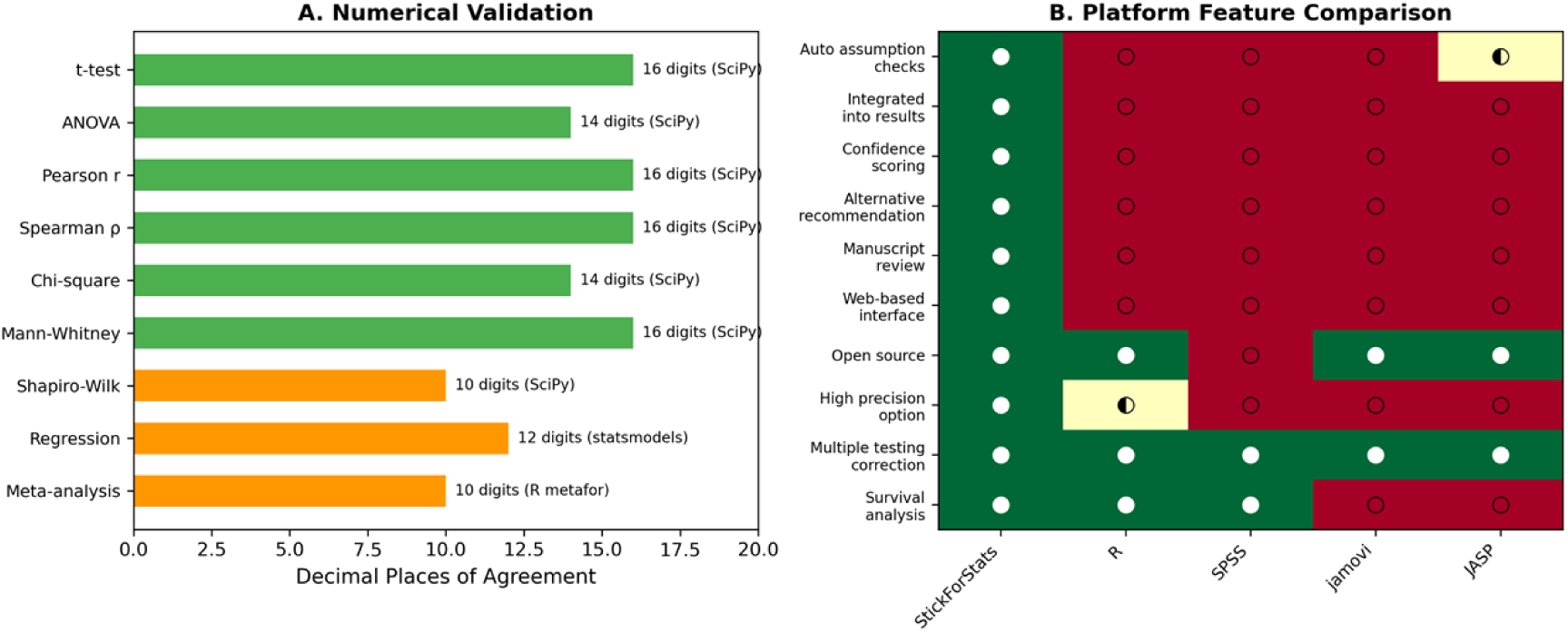
Platform comparison and validation. (A) Numerical agreement between StickForStats and reference implementations (SciPy, R, statsmodels) across 9 statistical test categories, demonstrating 10–16 decimal places of agreement. (B) Feature comparison heatmap of StickForStats vs. R, SPSS, jamovi, and JASP on 10 capabilities relevant to assumption validation and biomedical research.

### Limitations

We acknowledge several limitations. *Threshold dependence:* Guardian’s severity classifications depend on fixed thresholds (e.g., p < 0.05 for warnings); Guardian mitigates this by reporting actual test statistics, not just classifications. *Power of assumption tests:* Small samples may miss real violations while large samples may flag trivial ones; Guardian considers sample size in severity classification. *Incomplete coverage:* Guardian’s eight validators do not cover all possible assumptions—measurement reliability and selection bias may go undetected, though Guardian explicitly states which assumptions are checked. *Expert Mode override:* Experienced statisticians can proceed despite critical violations, though warnings remain visible. *Retrospective verification scope:* the consistency checker recomputes p-values from the reported statistic and degrees of freedom, so—like statcheck—it cannot recover a legitimately reported sphericity-corrected (Greenhouse-Geisser/Huynh-Feldt) p-value, cannot reproduce a multiplicity-adjusted (e.g. Tukey/Dunnett) p when only the raw statistic is shown, and can mistake a critical value used in a sample-size formula for a test result. In our 20-article evaluation these explainable cases accounted for 15 of the 19 flags (Table 7); flagged items are therefore recompute-versus-reported discrepancies for human review, not confirmed errors, and only four were genuine candidate reporting inconsistencies.

### Future directions

The platform includes five curated biological example datasets (CRISPR editing strategies, clinical trial survival, gene expression, epidemiological case-control, and dose-response) with documented analysis vignettes (availability details are given in Materials and Methods). Future work will expand the Bayesian analysis suite, add dose-response modeling for pharmacological studies, CONSORT flow diagram generation, and integrate with biological data repositories (GEO, ClinicalTrials.gov).

## Materials and methods

### Software implementation

The backend is implemented in Python 3.11 with Django 4.2 and Django REST Framework 3.14. Statistical computations use NumPy 1.25 [38] for array operations, SciPy 1.11 [39] for statistical functions, statsmodels 0.14 [40] for regression diagnostics and GLMs, lifelines 0.27 for survival analysis, and scikit-learn 1.3 for machine learning utilities. An optional high-precision mode uses mpmath 1.3 [27] for 50-decimal-digit calculations, critical for validation studies and extreme-value computations where IEEE 754 double precision (approximately 15 significant digits) may be insufficient. The frontend uses React 18 with Material-UI 5 for the user interface, Recharts 2.8 for interactive visualizations including forest plots and volcano plots, and jStat 1.9 for client-side computations. Asynchronous processing uses Celery 5.3 with Redis 7 for long-running analyses without blocking the interface. The platform is containerized with Docker and deployed via Docker Compose with PostgreSQL 15 for relational storage, Redis for caching and task queues, Nginx for static file serving, and optional Prometheus/Grafana monitoring. Guardian’s assumption checks are surfaced directly in the analysis interface alongside every result, making validation a visible default rather than an optional diagnostic (Fig 7).

**Fig 7.**
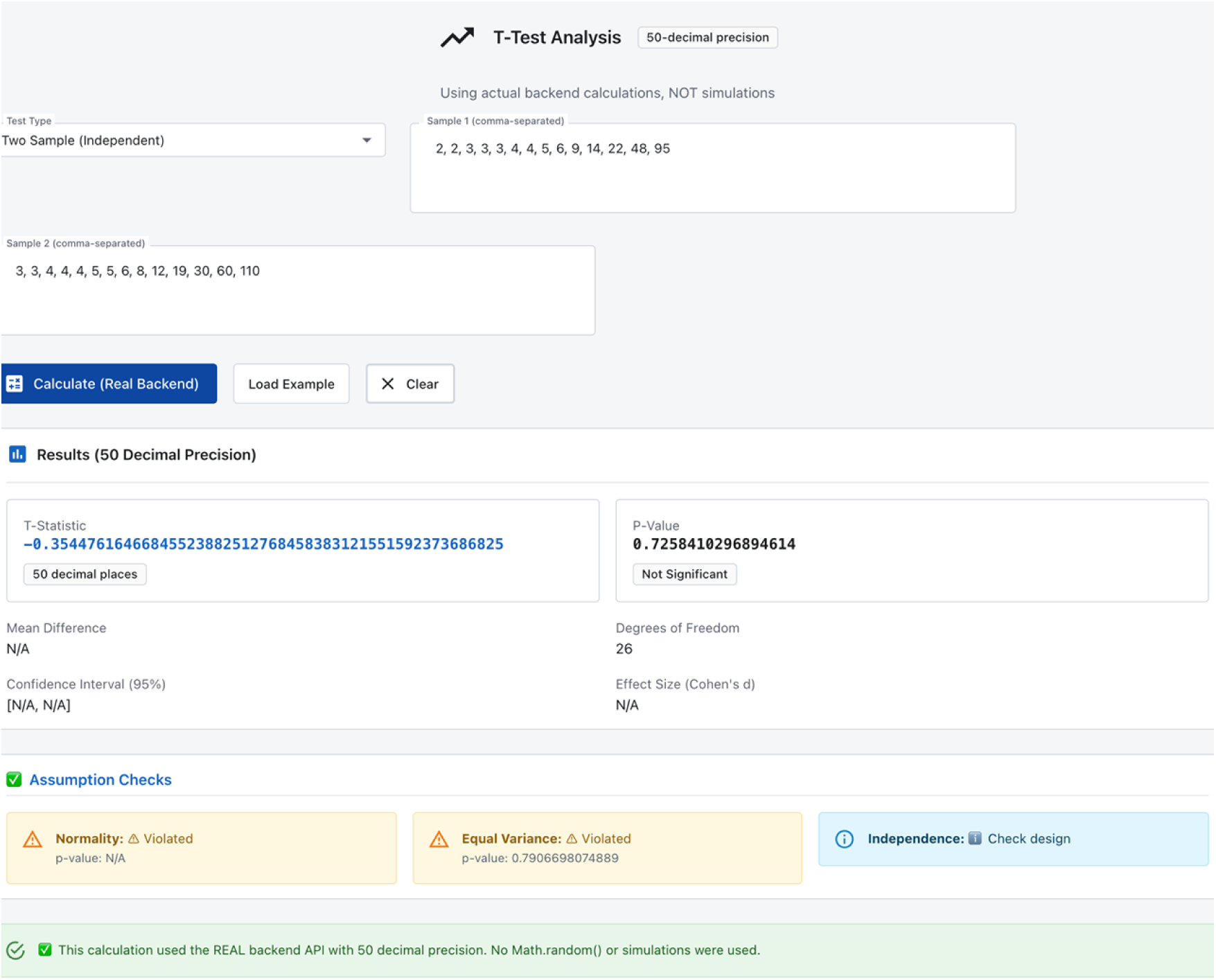
Guardian assumption validation in the StickForStats web interface. A two-sample t-test computed by the real backend at 50-decimal precision on a hosted instance of the platform. Before reporting, Guardian’s “Assumption Checks” panel evaluates each assumption and flags violations in place—here normality and equal variance are flagged as violated for the non-normal input, while independence is referred to study design—so the user sees the assumption status as an integral part of every analysis.

### Genomics differential expression workflow

The genomics module performs per-gene differential expression analysis with Guardian assumption validation. For each gene in an uploaded expression matrix, the service checks normality (Shapiro-Wilk) and variance homogeneity (Levene’s test) independently. Genes passing both checks are tested with the independent t-test (two groups) or ANOVA (multiple groups); genes failing either check are automatically cascaded to Mann-Whitney U or Kruskal-Wallis. After all per-gene tests complete, Benjamini-Hochberg FDR correction is applied across all raw p-values. The module generates volcano plot data (log2 fold change vs. negative log10 adjusted p-value) for visualization. In testing with log-normal gene expression data (100 genes, 20 samples), Guardian cascaded 98% of genes to nonparametric tests due to normality violations—the expected behavior for typical expression data.

### Validation methodology

Reference calculations were performed independently in R 4.3.2 and Python (SciPy 1.11) for all statistical tests. For each test type, we computed results on standard reference datasets and compared values to the maximum available floating-point precision. Meta-analysis results were cross-validated against R’s metafor package (DerSimonian-Laird estimator) to four-plus decimal places (see paper/replication/verify_meta_analysis_real.py for the 11-check verifier). G*Power cross-validation of power-analysis output is planned but not yet wired in this release; the in-app validation toggle reports “not implemented” rather than a fake match, and the manuscript’s validation summary therefore omits the power-analysis row. Seven Python scripts and one R script independently verify all reported values. Reproducibility scripts are provided in the repository under paper/replication/. For the retrospective-verification evaluation, the 20-article corpus was assembled from the PubMed Central open-access subset via a fixed E-utilities query (recorded in paper/replication/manuscript_validation/manifest.json); fetch_corpus.py rebuilds it and the validate_corpus management command reproduces the per-claim results (results.json). Each recomputable claim’s p-value was recomputed with SciPy and compared rounding- and inequality-aware against the reported value.

### Guardian evaluation

Each Guardian validator was validated against known datasets with confirmed properties: exponential distributions for normality (Shapiro-Wilk W = 0.886, p < 0.001, exact SciPy agreement), unequal-variance groups for homogeneity (Levene F = 8.92, p = 0.004, exact agreement), quadratic relationships for linearity (R-squared improvement = 45%, exact agreement with manual calculation). Edge case testing verified correct handling of empty arrays, single observations, identical values (zero variance), very large datasets (n = 10^6^, completed within 5 seconds), and extreme values (10^308^, no overflow errors).

## Availability and reproducibility

StickForStats is open-source under the MIT license; the version described here (v1.0.0, release tag v1.0.0) is openly available at https://github.com/visvikbharti/stickforstats_new, and a versioned snapshot will be archived on Zenodo to provide a citable DOI for the published version. The platform is platform-independent (Docker / Docker Compose) and runs in any modern browser; the backend requires Python >= 3.10, Django 4.2, PostgreSQL 15, and Redis 7 (SciPy >= 1.11, NumPy >= 1.24, statsmodels 0.14, mpmath 1.3). A Dockerfile and docker-compose.yml provision the full stack. A Python client SDK and command-line interface are available on PyPI (pip install stickforstats, or pip install stickforstats[cli] for the sfs command); the SDK connects to a StickForStats backend—a local Docker deployment or a hosted instance—through its REST API. All datasets analysed in this article are public and previously published; the replication package (paper/replication/) contains the verification scripts, the master runner (MASTER_VERIFICATION.py), and the statcheck head-to-head (statcheck_baseline.R) with run instructions. Per-dataset sources are: Fisher’s Iris (via scikit-learn); UCI Wine Quality (https://archive.ics.uci.edu/dataset/186/wine+quality); the IV-magnesium meta-analysis (Egger 1997 [26]; metafor::dat.egger2001 [33]); synovial-sarcoma RNA-seq (NCBI GEO accession GSE271517; Chen et al. 2024 [34]); and the 20-article retrospective-verification corpus (PubMed Central open-access subset, rebuilt by fetch_corpus.py from the recorded E-utilities query and manifest).

## Use of AI-assisted technologies

Generative AI (Claude, Anthropic) was used to assist both software development and the drafting of this manuscript. In software development, all AI-suggested code was reviewed by the authors, tested against reference implementations (SciPy, R), and validated through the project’s continuous integration pipeline (more than 1,500 automated tests across backend and frontend, all required checks green). In manuscript preparation, AI was used for drafting and editing assistance; all text was reviewed and verified by the authors, and every statistical value was independently recomputed and checked against SciPy and R. No AI tool is listed as an author. The authors take full responsibility for the content, accuracy, and integrity of this work, including any portion produced with AI assistance.

## Acknowledgments

We thank CSIR-Institute of Genomics and Integrative Biology for infrastructure and administrative support. We thank the developers of NumPy [38], SciPy [39], statsmodels [40], lifelines, and mpmath [27] whose libraries form the computational foundation of StickForStats.

## Supporting information

**S1 Text. Supplementary Information.** A single supporting document with five sections, all reproducible from the open-source repository (https://github.com/visvikbharti/stickforstats_new): (S1.1) Guardian validator specifications — the eight validators, the assumption each checks, its statistical method, and the test-type-to-validator mapping; (S1.2) Programmatic access — Python SDK (pip install stickforstats) and sfs CLI usage examples; (S1.3) Additional validation on standard R datasets — Guardian results on mtcars (regression), ToothGrowth (two-sample t-test), and PlantGrowth (one-way ANOVA), reproducible via paper/replication/additional_real_data_analysis.py; (S1.4) Guardian test-suite coverage — per-suite test counts (22 integration, 16 middleware, 46 validator-unit, 12 math-correctness backend tests; 25 component and 30 hook frontend tests); (S1.5) Performance benchmarks — end-to-end API latency with and without the Guardian pipeline (mean ± SD over 100 requests; paper/replication/benchmark_api.py), showing a median Guardian overhead of 0.2 ms with all latencies below 10 ms at the 99th percentile.

## References

1. Baker M. 1,500 Scientists Lift the Lid on Reproducibility. Nature. 2016;533(7604):452–454.

2. Open Science Collaboration. Estimating the Reproducibility of Psychological Science. Science. 2015;349(6251):aac4716.

3. Ioannidis JPA. Why Most Published Research Findings Are False. PLoS Medicine. 2005;2(8):e124.

4. Zimmerman DW. A Note on Preliminary Tests of Equality of Variances. Br J Math Stat Psychol. 2004;57(1):173–181.

5. Zimmerman DW. Comparative Power of Student t Test and Mann-Whitney U Test. J Exp Educ. 2004;73(2):167–183.

6. Hoekstra R, Kiers HAL, Johnson A. Are Assumptions of Well-Known Statistical Techniques Checked, and Why (Not)? Front Psychol. 2012;3:137.

7. Keselman HJ, et al. Statistical Practices of Educational Researchers. Rev Educ Res. 1998;68(3):350–386.

8. Osborne JW. Improving Your Data Transformations: Applying the Box-Cox Transformation. Pract Assess Res Eval. 2010;15(12):1–9.

9. Benjamini Y, Hochberg Y. Controlling the False Discovery Rate. J R Stat Soc B. 1995;57(1):289–300.

10. Schulz KF, Altman DG, Moher D. CONSORT 2010 Statement. BMJ. 2010;340:c332.

11. DerSimonian R, Laird N. Meta-Analysis in Clinical Trials. Control Clin Trials. 1986;7(3):177–188.

12. Nickerson RS. Confirmation Bias: A Ubiquitous Phenomenon. Rev Gen Psychol. 1998;2(2):175–220.

13. Appelbaum M, et al. Journal Article Reporting Standards for Quantitative Research in Psychology. Am Psychol. 2018;73(1):3–25.

14. Nosek BA, Ebersole CR, DeHaven AC, Mellor DT. The Preregistration Revolution. Proc Natl Acad Sci. 2018;115(11):2600–2606.

15. Nuijten MB, Hartgerink CHJ, van Assen MALM, Epskamp S, Wicherts JM. The prevalence of statistical reporting errors in psychology (1985-2013). Behav Res Methods. 2016;48(4):1205–1226.

16. Aust F, Barth M. papaja: Prepare Reproducible APA Journal Articles with R Markdown. R package version 0.1.0.9997. 2020.

17. Shapiro SS, Wilk MB. An Analysis of Variance Test for Normality. Biometrika. 1965;52(3-4):591–611.

18. Anderson TW, Darling DA. A Test of Goodness of Fit. J Am Stat Assoc. 1954;49(268):765–769.

19. Levene H. Robust Tests for Equality of Variances. In: Contributions to Probability and Statistics. Stanford University Press; 1960:278–292.

20. Brown MB, Forsythe AB. Robust Tests for the Equality of Variances. J Am Stat Assoc. 1974;69(346):364–367.

21. Durbin J, Watson GS. Testing for Serial Correlation in Least Squares Regression. II. Biometrika. 1951;38(1/2):159–177.

22. Grubbs FE. Procedures for Detecting Outlying Observations in Samples. Technometrics. 1969;11(1):1–21.

23. Cohen J. Statistical Power Analysis for the Behavioral Sciences. 2nd ed. Hillsdale, NJ: Lawrence Erlbaum; 1988.

24. Wald A, Wolfowitz J. On a Test Whether Two Samples Are from the Same Population. Ann Math Stat. 1940;11(2):147–162.

25. Breusch TS, Pagan AR. A Simple Test for Heteroscedasticity. Econometrica. 1979;47(5):1287–1294.

26. Egger M, et al. Bias in Meta-Analysis Detected by a Simple, Graphical Test. BMJ. 1997;315(7109):629–634.

27. Johansson F. mpmath: A Python Library for Arbitrary-Precision Floating-Point Arithmetic. 2013.

28. Faul F, et al. G*Power 3: A Flexible Statistical Power Analysis Program. Behav Res Methods. 2007;39(2):175–191.

29. Fisher RA. The Use of Multiple Measurements in Taxonomic Problems. Ann Eugen. 1936;7(2):179–188.

30. Cortez P, et al. Modeling Wine Preferences by Data Mining. Decis Support Syst. 2009;47(4):547–553.

31. Bharti V, Chakraborty D. CRISPRArchitect v3: Multi-nuclease, consequence-guided decision support for genome editing strategy design. GitHub. 2026. https://github.com/visvikbharti/CRISPRArchitect

32. Sterne JAC, Egger M. Funnel Plots for Detecting Bias in Meta-Analysis: Guidelines on Choice of Axis. J Clin Epidemiol. 2001;54(10):1046–1055.

33. Viechtbauer W. Conducting Meta-Analyses in R with the metafor Package. J Stat Softw. 2010;36(3):1–48.

34. Chen Y, Su Y, Cao X, Siavelis I, Leo IR, Zeng J, Tsagkozis P, Hesla AC, Papakonstantinou A, Liu X, Huang W-K, Zhao B, Haglund C, Ehnman M, Johansson H, Lin Y, Lehtiö J, Zhang Y, Larsson O, Li X, de Flon FH. Molecular Profiling Defines Three Subtypes of Synovial Sarcoma. Adv Sci (Weinh). 2024;11(41):e2404510. doi:10.1002/advs.202404510. PMID: 39257029. PMCID: PMC11892499.

35. R Core Team. R: A Language and Environment for Statistical Computing. Vienna, Austria: R Foundation; 2023.

36. JASP Team. JASP (Version 0.17.3). 2023.

37. The jamovi project. jamovi (Version 2.4). 2023.

38. Harris CR, et al. Array Programming with NumPy. Nature. 2020;585(7825):357–362.

39. Virtanen P, et al. SciPy 1.0: Fundamental Algorithms for Scientific Computing in Python. Nat Methods. 2020;17(3):261–272.

40. Seabold S, Perktold J. Statsmodels: Econometric and Statistical Modeling with Python. Proc 9th Python Sci Conf. 2010:57–61.

